# Dopamine alters motor learning performance in the presence and absence of feedback

**DOI:** 10.1101/2024.11.23.625034

**Authors:** Li-Ann Leow, Ashley Huey-Ryu Tan, Timothy J Carroll, Rob Adam, Paul E Dux, Hannah L Filmer

## Abstract

We often choose to learn motor skills not only because of some external reward, but also because motor learning, in and of itself, is satisfying. While dopamine is thought to drive reward-based motor learning, it remains unclear whether dopamine is implicated in motor learning under conditions ostensibly driven by intrinsic rewards/motivation (i.e., in the absence of extrinsic feedback or reward). Here, by pharmacologically manipulating dopamine using the dopamine precursor Levodopa, we investigated the role of dopamine in an explicit motor learning task guided by internally determined signals of performance success, by removing intrinsic feedback about task success. Specifically, we asked participants to strategically aim away from presented targets by various instructed angles: a form of explicit motor learning that contributes to performance in the classic visuomotor rotation task. In the feedback condition, targets jumped mid-movement by the instructed angle, such that success in target hitting depended on successfully aiming away by the instructed angle. In the no-feedback condition, intrinsic feedback about task success was removed by having targets disappear mid-movement, such that participants could not know if they succeeded at hitting the targets or not. We found that dopamine altered performance, both with and without task feedback about task success. This provides direct evidence for a role of dopamine in motor learning driven by internal task goals.

---

Motor learning is often driven by extrinsic reward: to play her favourite app-based game, a child quickly learns the sequence of finger-swipes needed to unlock her parent’s mobile phone. Extrinsic rewards can take the form of primary (e.g., food, water) or secondary rewards (e.g., points or money) contingent upon achieving some performance criteria. The promise of reward not only drives movements to be more vigorous (e.g., Carpenter & Williams, 1995; Milstein & Dorris, 2007; Manohar *et al*., 2015; Reuter *et al*., 2018; Summerside *et al*., 2018; Carroll *et al*., 2019; Galaro *et al*., 2019), but can also improve the acquisition (Nikooyan & Ahmed, 2015) and retention (Abe *et al*., 2011; Galea *et al*., 2015; Dayan *et al*., 2016; Hamel *et al*., 2019; Spampinato *et al*., 2019; van der Kooij & Smeets, 2019; Sporn *et al*., 2022; Yin *et al*., 2023) of motor learning. A key mechanism through which reward alters behaviour is via reward prediction errors, defined as discrepancies between predicted and actual rewards (Sutton & Barto, 1998). Reward prediction errors trigger phasic firing of midbrain dopamine neurons (Schultz *et al*., 1997), which alters the vigour of movements (Sedaghat-Nejad *et al*., 2019) and shapes motor behaviour (Izawa & Shadmehr, 2011; Cashaback *et al*., 2017).

Yet we also often learn for its own sake. In the absence of extrinsic rewards, a child learns how to ride a scooter not because she receives some reward, but because scooter-riding is inherently fun. The distinction between behaviour driven by intrinsic motivation and extrinsic rewards has long been recognised (Harlow, 1950; Ryan & Deci, 2000; Baldassarre & Mirolli, 2013). Although we do not fully understand the drivers of intrinsic motivation, it appears to be modulated by internal rewards associated with meeting task goals (Blain & Sharot, 2021). Information about task success might thus modulate intrinsic motivation. In experimental contexts, information about task success can alter motor learning (Schaefer *et al*., 2012; Cashaback *et al*., 2017; Leow *et al*., 2018; Palidis *et al*., 2018; Sidarta *et al*., 2018; Kim *et al*., 2019; Leow *et al*., 2020; Sadaphal *et al*., 2022; Tsay *et al*., 2022), and its effects seem separable to extrinsic rewards contingent upon task performance (Vassiliadis *et al*., 2021).

How does the brain engage in motor learning in the absence of extrinsic rewards? Whilst an increasing body of work characterises the neural mechanisms of learning driven by extrinsic rewards, comparatively little is known about how we learn and act in the absence of extrinsic rewards (Blain & Sharot, 2021). Work in animal models implicates the dopamine system. For example, dopamine neurons in zebra finches show spontaneous activity that correlates with song performance, even in the absence of extrinsic rewards, cues, or external perturbations (Duffy *et al*., 2022). In humans, learning and performing actions in the absence of extrinsic rewards engages midbrain dopamine-dependent reward networks that are also involved in processing primary rewards, such as the ventral striatum (Ripolles *et al*., 2016) and the ventromedial prefrontal cortex (Chew *et al*., 2021). This supports the view that intrinsic rewards alters behaviour mostly by co-opting existing dopamine-dependent neural circuitry used to process primary rewards (Blain & Sharot, 2021). However, intrinsic and extrinsic rewards evoke subtly different brain responses: for example, learning with and without extrinsic rewards seems to differentially implicate the dorsal and ventral striatum (Lutz *et al*., 2012; Ripolles *et al*., 2016), areas with substantial functional heterogeneity (Collins & Saunders, 2020). It thus remains unclear if and how processing of intrinsic and extrinsic rewards might differentially implicate brain dopamine function during motor learning.

Here, in a pre-registered study, we asked the following question: how does dopamine contribute to motor learning driven by internal goals in the absence of extrinsic rewards? We asked participants to complete an aiming task, where they were explicitly instructed to aim away from presented targets by angles that ranged in size, where larger aiming angles are more difficult. This task is also termed the visuomotor mental rotation task (Georgopoulos & Massey, 1987; Neely & Heath, 2010). Such learning to aim away from a presented target is a key explicit strategy used to counteract rotated cursor feedback in the classic visuomotor rotation task (Leow *et al*., 2017; McDougle & Taylor, 2019), which can drive performance improvements in these tasks (Leow *et al*., 2020). In this instructed aiming task, participants were instructed to hit as many targets as possible. Participants were either deprived of information about the task success (targets always disappeared mid-movement, thus precluding knowledge of target-hits/misses)(Taylor & Ivry, 2011); or given information about task success (targets jumped mid-movement by the instructed aiming angle, such that participants were rewarded with target hits and avoided target-misses when they succeeded at the task of aiming by the instructed angle). Such feedback about task success appears important to the explicit, strategic component of sensorimotor adaptation, as it often alters accuracy when the task goal is to reach to the target (Leow *et al*., 2020; Sadaphal *et al*., 2022; Tsay *et al*., 2022), and thus we expected this task feedback to also increase aiming accuracy and/or shorten reaction times. Task success information might directly affect motor performance, and/or affect motor performance indirectly, by altering intrinsic motivation to attain task goals. We assessed the role of dopamine in learning and performance in the absence or presence of task success information, by having participants consume Levodopa, a dopamine precursor which increases brain dopamine availability. We hypothesized that if dopamine contributes to performance by primarily altering the processing of extrinsic task success information, then dopamine will only alter motor performance in the presence of feedback about task success. Alternatively, if dopamine alters performance primarily by modulating intrinsic motivation, then manipulating dopamine will affect performance in a similar fashion regardless of the absence/presence of feedback about task success. Finally, if dopamine alters learning regulated by internal estimations of task performance and learning regulated by extrinsic task success information, then dopamine will alter performance both in the presence and absence of feedback about task success, but effects will differ between conditions.

## 2. Method

### 2.1. Participants

Participants were recruited from The University of Queensland community and were reimbursed for participation (AUD$20/hour). Participants were screened for neurological and psychiatric conditions, contraindications for Levodopa, and injury preventing the use of their right-arm and provided written informed consent. In accordance with the National Health and Medical Research Council’s guidelines, this experiment was approved by the human ethics committee at The University of Queensland. No datasets were excluded from the study.

To the best of our knowledge, there are no equivalent studies examining how dopamine alters motor performance on a visuomotor mental rotation task in the presence or absence of extrinsically defined task errors. We used a Bayesian stopping rule to optimize sample sizes, as stated in our preregistration (https://osf.io/m7a3g/). Specifically, we intended to stop data collection either upon obtaining moderate evidence for including or excluding an effect of dopamine (BF_incl_ > 3 or BF_excl_ > 3) for aiming accuracy or reaction times, or upon collecting 80 complete datasets (20 participants for each of the 4 conditions), in line with previous studies which examine effects of Levodopa on motor performance in neurotypical young adults. We reached our stopping rule of BF_incl_ > 3 upon collecting the minimum of 15 participants per group.

### 2.1. Study design

In this 2 x 2 between-subjects design manipulating drug (placebo, levodopa) and feedback about task success (no feedback, feedback), participants were randomly assigned to complete one of the following four study conditions: (1) placebo + feedback (n=15), (2) levodopa + feedback (n=17), (3) placebo + no-feedback (n=15), and (4) levodopa + no-feedback (n=16). Randomisation of study conditions to each participant was performed using https://www.random.org/. The feedback manipulation is detailed later. Briefly, in the task feedback condition, targets “jumped” mid-movement by the instructed aiming angle, such that target-hits were contingent upon successful aiming. In the no-feedback condition, targets disappeared mid-movement, removing the possibility of target hits regardless of aiming success: participants thus had no feedback about their task success.

### 2.3. Drug manipulation

Participants ingested either Madopar 125 tablet (100mg Levodopa and 28.5mg benserazide hydrochloride) or a vitamin C tablet (Placebo). Tablets were crushed and dispersed in orange juice to achieve participant blinding. Drug administration was completed by an experimenter uninvolved in data collection, out of view of the experimenter responsible for data collection, to achieve experimenter blinding. This method is effective for double-blinding (Beierholm *et al*., 2013; Leow *et al*., 2023; Leow *et al*., 2024). We chose Levodopa as it increases brain dopamine levels without affecting norepinephrine levels (Everett & Borcherding, 1970), and achieves peak plasma levels within approximately 45 minutes of ingestion. Participants waited 45 minutes after drug ingestion to complete the aiming task. During this wait period, we assessed participants trait impulsivity by asking participants to complete the Barratt Impulsivity Scale, and assessed working memory capacity by asking participants to complete either the memory updating task or the operation span task (Lewandowsky *et al*., 2010); the other task was completed at the end of the session). These measures were taken as proxy measures of individual baseline dopamine function, which has previously been found to be partly predicted by impulsivity (Buckholtz *et al*., 2010) and working memory (Cools *et al*., 2008). Some studies find such measures can partly predict effects of pharmacological manipulations of dopamine (Clatworthy *et al*., 2009; Broadway *et al*., 2018; Fallon *et al*., 2019): thus, we intended to include these measures as covariates of no interest in our analyses.

To assess for changes in arousal, heart rate, blood pressure, and mood were measured at three timepoints (1) before drug ingestion, (2) approx. 15-20 minutes post-ingestion, and (3) post-study completion (approx. 150 minutes post-ingestion). Mood was assessed using the Mood Rating Scale (Bond & Lader, 1974) which includes sixteen items separated in following factors: alertness, contentedness and calmness.

### 2.4. Apparatus

We used a vBot planar robotic manipulandum (Howard *et al*., 2009), which has a low-mass, two-link carbon fiber arm and measures position with optical encoders sampled at 1,000 Hz. Participants completed the aiming task while seated on a height-adjustable chair at their ideal height for viewing the screen. Visual feedback was presented on a horizontal plane on a 27” LCD computer monitor (ASUS, VG278H, set at 60-Hz refresh rate) mounted above the vBOT and projected to the subject via a mirror in a darkened room. Participants had no direct vision of their hand. Hand position was represented by a red cursor (0.25 cm radius circle) on a black background. The mirror also allowed visual feedback of the target (a 0.5 cm-radius circle) and the start location (a 0.35 cm radius circle), which was aligned 10 cm to the right of the participant’s midsagittal plane at approximately mid-sternum level.

Participants also completed a control block (0°) where they reached straight towards the target (not depicted), as well as an initial familiarisation block 90° to acquaint them with the task conditions (not depicted). **E.** Participants completed sets of blocks for each of the aiming angles. In each set, participants completed an Instruction block (24 trials) with aiming landmarks to teach participants to aim away by the instructed angle, a Training block without landmarks (80 trials), but with the task feedback manipulation, a no-feedback Probe block (40 trials) where participants continued to aim by the instructed angle, a Washout block (40 trials) where participants stopped aiming by the instructed angle. To economise on session length, there was no Instruction, Probe, or Washout blocks for the 0° set. The order of the aiming angles was the same for all participants: 90° (familiarisation), 130°, 30°, 0°, 150°, 60°, 160°, 120°.

### 2.5. Aiming task

#### 2.5.1. Trial structure

The design of our aiming task was similar to previous visuomotor mental rotation studies (Georgopoulos & Massey, 1987; Pellizzer & Georgopoulos, 1993; Neely & Heath, 2010). In the aiming task, participants were asked to aim towards targets by an instructed aiming angle. At the start of each trial, participants saw a grey start position (0.35cm radius circle) aligned approximately 10cm to the right of participant’s midsagittal plane, mid-sternum level, and a red virtual circular cursor (a 0.25cm radius circle) representing robot handle position against a black background. With their right hand holding the robot handle, participants first positioned the cursor in the start circle. Once the cursor stayed within the start circle for 2 seconds, a yellow target (0.5cm radius circle), accompanied by an auditory signal, appeared at one of eight possible target directions, in random order (0°, 45°, 90°, 135°, 180°, 225°, 270°, or 315°), 9cm away from the start. Target directions were pseudorandomised such that each of the 8 possible targets directions occurred once every 8 trials (every 8 trials was 1 bin). Once a trial was initiated, if no initiation of arm movement was detected within 6 seconds, the trial was terminated. Similarly, in instances where the participant failed to achieve the required movement extent (the 9cm start-target distance), the trial was terminated. Terminated trials were replaced at the end of each 8-trial bin.

#### 2.5.2. Instructions and Block structure

Participants were first provided with the following set of instructions, which emphasized aiming accuracy “You are about to perform sets of trials in which you will be asked to move at specific angle relative to the target in clockwise direction. The aiming angle will differ between blocks. We will tell you what the angle is for each block. First, make up your mind and then try to keep your movement straight and fast.

Try your best to execute your movements as accurately as possible and to hit as many targets as possible. Between each set we will ask you to aim at a 0-degree angle, that means you should hit the target directly. These sets will contain 40 trials.”

Next, participants were asked to complete sets of trials for each of the aiming angles (see Figure 1E). At the start of each set, they were first completed an Instruction block (3 bins, or 24 trials) where they were provided with a ring of explicit aiming landmarks (letters and numbers) displayed around the target positions, and participants were asked to aim directly towards the letter ‘A’ which was positioned at the instructed aiming angle relative to the current target, similar to Taylor *et al*. (2014).

**Figure 1.**
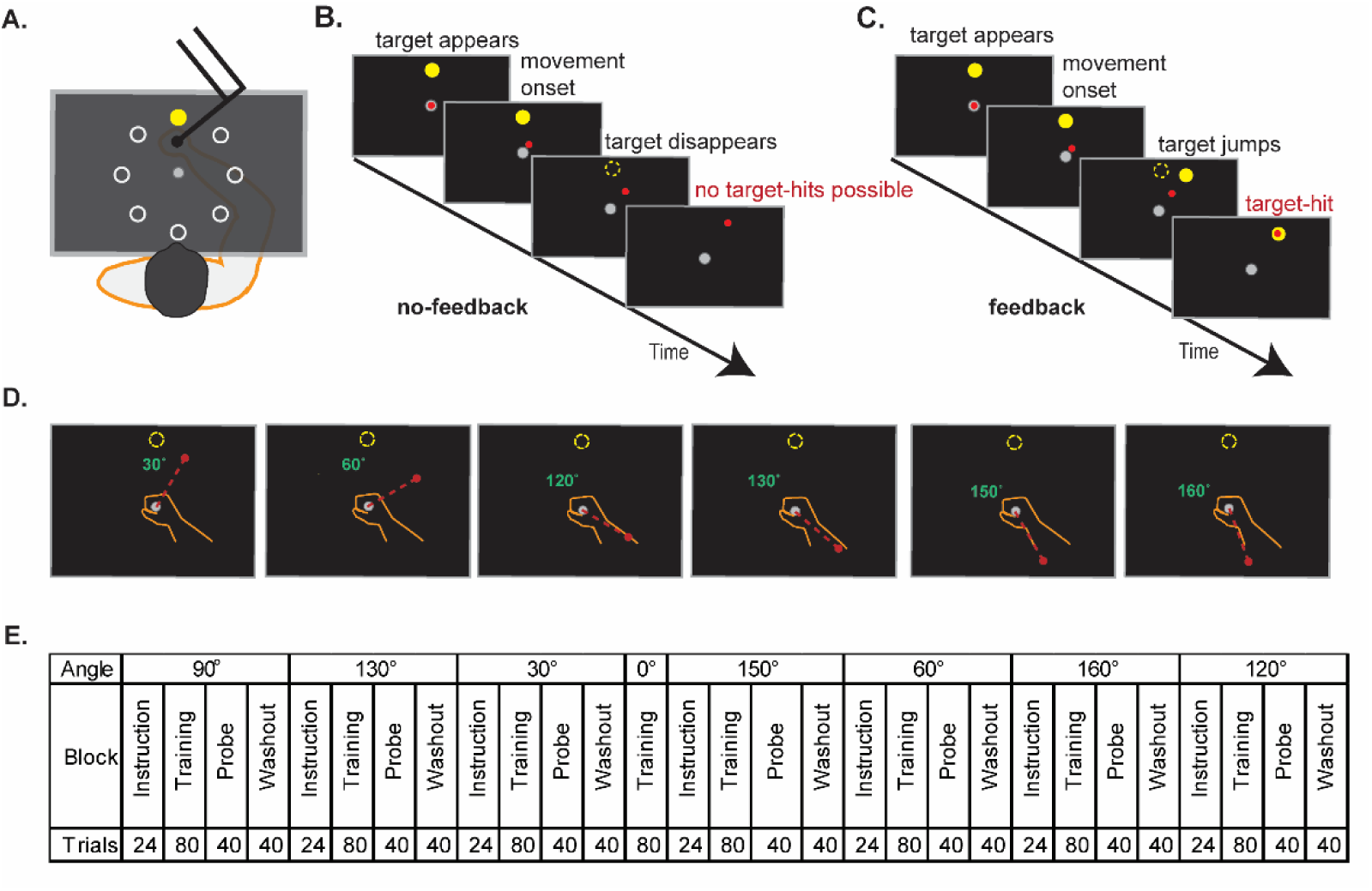
**A**. Participants completed the aiming task by aiming away from one of eight possible targets (each 1 of the 8 target positions appeared in random order in every bin, where 1 bin= 1 visit to each of the 8 targets). This task resembles those of (Georgopoulos & Massey, 1987; Neely & Heath, 2010) **B**. In the no-feedback condition, participants did not know whether they successfully hit the target, as the targets would disappear mid-movement, making target-hits impossible. **C**. In the feedback condition, as the target would “jump” mid-movement by the instructed aiming angle, successful aiming by the instructed aiming angle would result in target hits. **D**. Participants completed sets of trials where they had to aim away from the presented targets by the instructed angles (30°, 60°, 120°, 130°, 150°, 160°), clockwise.

This was followed by a Training block (80 trials, or 10 bins), where participants had to aim by the same instructed angle as in the Instruction block. During training, participants either received feedback about task success/task failure, or received no feedback, depending on assigned condition. In the feedback condition, feedback on task success was given by having the target “jump” mid-movement by the instructed aiming angle: successful aiming resulted in a target-hit, unsuccessful aiming resulted in a target miss. This “jump” was achieved by extinguishing the target once the movement extent exceeds 4cm (start-target distance = 9cm), and immediately re-displaying the target at the ideal aiming direction. In the no-feedback condition, targets were extinguished once the movement extent exceeded 4cm: preventing the experience of target hits or target misses.

After the training block, participants completed a Probe block (40 trials, or 5 bins) where they were asked to continue to aim by the same instructed angle, but now without task success feedback across all participants (i.e., targets disappeared mid-movement).

Finally, participants completed Washout blocks (40 trials, or 5 bins) where they were instructed to aim directly towards the presented target, which remained visible throughout the trial.

In each set, participants aimed away from the presented target by a prespecified angle, clockwise. To familiarise participants with the task, participants completed a familiarisation set with a 90° aiming angle. Subsequently, all participants completed the following sets: 130°, 30°, 0°, 150°, 60°, 160°, 120°, in this order. To reduce task duration, we did not include the landmark, probe, or washout trials for the straight ahead (0°) aiming angle. Participants completed a total of 1368 trials, which was approximately 2 hours long.

### 2.6. Data analysis

Our analyses followed our preregistration analysis plan (https://osf.io/m7a3g/). As our task instructions emphasized accurate aiming, our primary dependent variable of interest was aiming accuracy, which was estimated from absolute directional error: the unsigned difference between the ideal aiming angle and the absolute values of the actual reach direction, where reach direction was measured at 20% of the start-target distance (i.e., 1.8 cm into the 9cm start-target distance). In standard visuomotor rotation tasks, the majority of reach directions reflect incomplete adaptation to the cursor rotation (and thus, for a clockwise rotation, most reach directions will be in the counterclockwise direction relative to the target). However, our aiming task (also termed the visuomotor mental rotation task) asks participants to aim away from the presented target by a specified angle without employing any cursor rotation. In these tasks, approximately half the reach directions tend to be in the expected direction (i.e., clockwise if asked to aim clockwise), and half tend to be in the other direction (Neely & Heath, 2010). This was also true for our data set: an averaged 49% of reach directions in the training trials in the expected direction, 51% of reach directions were in the opposite direction. Because of this, mean-averaging such signed directional data across bins for analysis and for graphical depiction in figures is problematic, as these would result in misleading mean values that are close to zero. Thus, we focussed on absolute unsigned errors in during re-aiming.

We considered absolute unsigned errors in excess of 90° to be outliers and discarded these from analyses of errors (0.40% of trials in the landmark blocks, 0.25% of trials in the training blocks, 0.49% of trials in the probe blocks, 0.07% trials in the washout blocks). We also assessed the number of target-hits (i.e., the number of trials where participant’s 0.25cm radius cursor was in contact with the 0.5cm radius target), as an index of success in achieving the task goal of hitting as many targets as possible. Note that we designed our cursor radius to be half that of our previous study (Leow *et al*., 2024), to ensure that the task feedback manipulations were sufficiently sensitive.

Whilst our instructions and task structure emphasised aiming accuracy, we were also interested in evaluating changes in the speed of movement planning. To this end, we also quantified reaction time, defined as the interval from target appearance to movement onset, where movement onset was the time at which the hand speed first exceeded 2 cm/s. In addition, as dopamine has been implicated in movement vigour, we additionally examined how the dopamine drug manipulation and the task feedback manipulation altered peak velocities in exploratory analyses.

In accordance to our preregistered analysis plan, to test the effect of the drug and target-jump manipulations on aiming performance, we ran Bayesian Repeated-Measures ANOVAs with within-subjects variables Angle (0°, 30°, 60°, 120°, 130°, 150°, 160°) and Bin (1….10) and between-subjects variables Drug (Placebo, Levodopa) and Condition (Target Jump, Target Disappear) on dependent variables of interest, for the instruction block, training block and the post-training probe block, and the washout block. To evaluate how the task feedback manipulations altered performance and training-related performance gains in the absence and presence of levodopa, we ran follow-up Feedback x Angle x Bin ANOVAs, separately for the placebo and levodopa groups. To evaluate how levodopa altered performance and training-related performance gains with and without task feedback, we ran follow-up Drug x Angle x Bin ANOVAs, separately for groups with and without task feedback. Statistical analyses were run using JASP 0.19.3. Random slopes were not included, as per (Morey & Rouder, 2022). We made inferences using Bayesian statistics, quantifying evidence for the test hypothesis, as well as evidence for the null hypothesis. Inclusion Bayes factors (BFinclusion, BF_incl_) were determined to estimate the strength of evidence in favour of including an effect relative to models stripped of that effect.

Exclusion Bayes factors (BFexclusion, BF_excl_) were determined to estimate the strength of evidence in favour of excluding an effect relative to models stripped of that effect. In post hoc tests, where evidence for the null hypothesis is reported as BF_01_, and evidence for the test hypothesis is reported as BF_10_, the posterior odds were corrected for multiple testing by fixing to 0.5 the prior probability that the null hypothesis holds across all comparisons (Westfall *et al*., 1997). We used Jeffreys’s evidence categories for interpretation of the reported Bayes factors (Wetzels *et al*., 2011). Specifically, Bayes factors of 1–3 were interpreted as anecdotal, 3–10 as moderate, and >10 as strong evidence for the test hypothesis, inclusion Bayes factors of 1–3 were interpreted as anecdotal, 3–10 as moderate, and >10 as strong evidence for the hypothesis.

In our preregistration, we stated that we would enter impulsivity and working memory capacity as covariates of no interest, following previous work that use such measures as a proxy of striatal dopamine synthesis capacity (Cools *et al*., 2008; Buckholtz *et al*., 2010), as they are thought to predict responsivity to dopamine drug manipulations (Cools *et al*., 2008; Clatworthy *et al*., 2009; Froböse *et al*., 2018; Fallon *et al*., 2019),. However, new evidence shows evidence against the use of impulsivity or working memory capacity as a proxy of striatal dopamine synthesis capacity (van den Bosch *et al*., 2023). Furthermore, because our Bayesian stopping rule stipulated that we stopped data collection upon showing moderate evidence for an effect of drug, we stopped data collection at a relatively small sample size that precludes strong inferences about these covariates. Results were not meaningfully altered by including the covariates (i.e., the effects observed were unchanged with or without including these covariates). We thus decided against entering impulsivity and working memory capacity as covariates in our analyses.

## 3. Results

### 3.1. Baseline checks and manipulation checks

In our within-subjects task design, participants completed multiple angle-sets, and the task feedback manipulation was applied in all training blocks for all angle-sets. Thus, the instruction blocks in each angle-set were not “pure” baselines where they had no prior experience of task feedback: only the instruction block for the 90° familiarisation set preceded any experience of task feedback. We thus ran analyses to check if the task feedback groups showed pre-existing differences in this instruction block for 90°. We did not find clear evidence for pre-existing differences between the task feedback groups. As participants familiarised themselves with the task, both reaction times and error reduced across bins (reaction time, BF_incl_ = 4.390E+17, error BF_incl_ =1044.194). There was anecdotal-moderate evidence for no existing baseline differences across feedback groups for reaction time (main effect of feedback, BF_excl_ = 2.229; bin x feedback interaction, BF_excl_ = 2.675, bin × drug × feedback interaction: BF_excl_ = 6.690) and for error (main effect of Feedback, BF_excl_ = 3.774, bin x feedback: BF_excl_ = 1.873, bin × drug × feedback: BF_excl_ = 1.343 Next, we conducted manipulation checks to test whether our explicit instructions and aiming landmarks, given in the pre-training instruction block, improved task performance. Then, we assessed whether performance improved across bins during the training block.

#### 3.1.1. Explicit task instructions and aiming landmarks improved performance

Prior to each training block, participants first completed an instruction block of 3 bins (24 trials) where they were given explicit task instructions and landmarks to learn to aim at the required aiming angles: the landmark A indicated where participants needed to aim towards. As participants first learnt how to aim by the instructed angles, reaction times were longer than the later training blocks, as shown in the gray regions in Figure 2. Reaction times reduced from the first instruction bin to subsequent instruction bins (main effect of bin for reaction time BF_incl_ = 4.70E+29).

**Figure 2.**
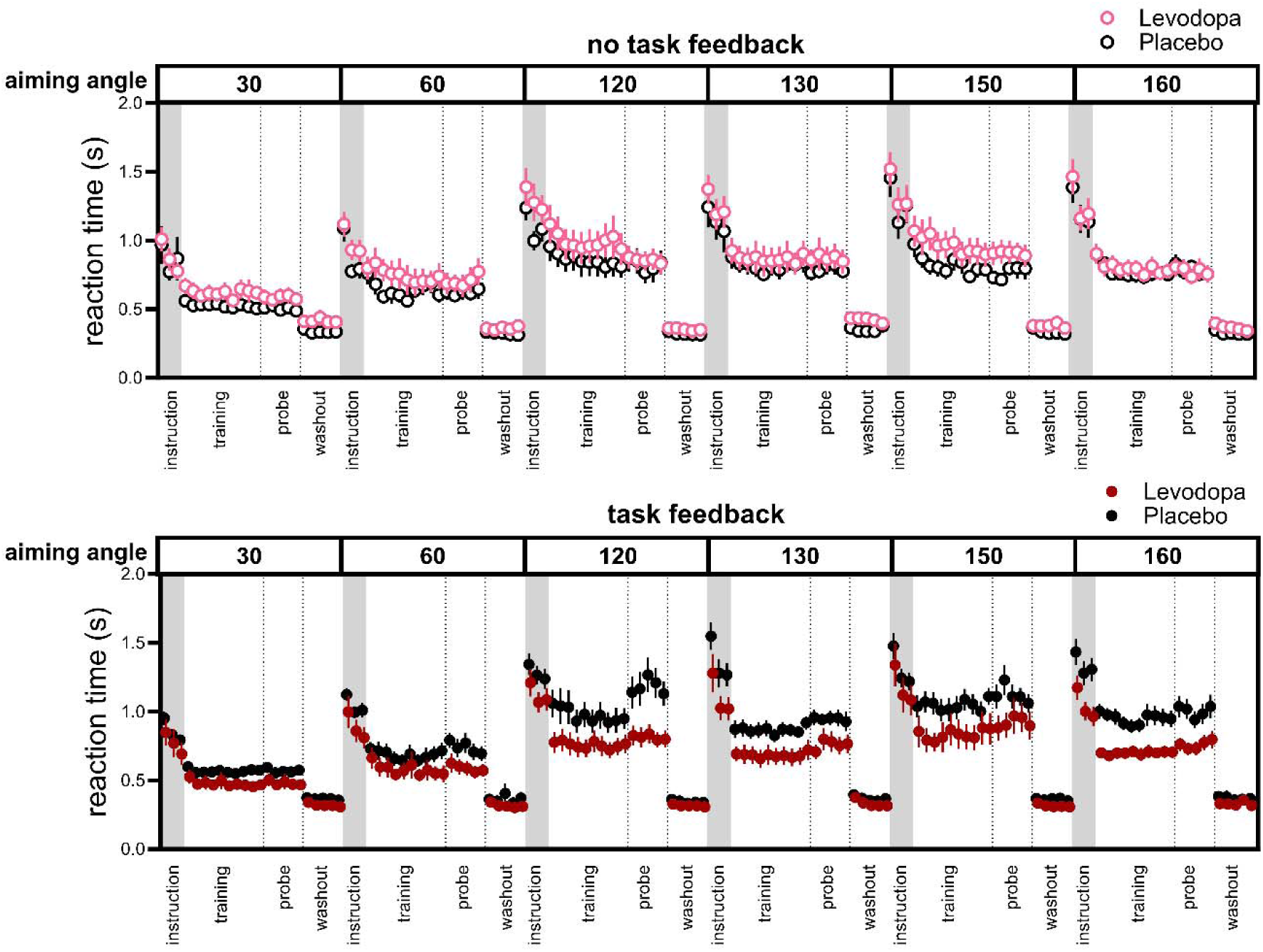
Bin-by-bin reaction times in each block, for each of the aiming angles, for participants who completed training without task feedback (top panel) and with task feedback (bottom panel). Reaction times were longer when first learning to re-aim in the instruction blocks, gradually reduced with training, and reduced further when participants were no-longer required to re-aim in the washout blocks.

In contrast to reaction times which reduced across instruction bins, accuracy did not reduce systematically across the instruction bins, possibly due to the presence of aiming landmarks (no evidence for main effect of bin for error, BF_incl_ = 0.504). The effect of Levodopa on reaction time and error in the instruction blocks was not clear. Whilst there was moderate evidence for drug x feedback interactions for error (BF_incl_ = 3.337), follow-up analyses failed to show evidence for an effect of Levodopa in these instruction blocks, both in participants who received task feedback, and in those deprived of task feedback in the training blocks (all Bayes factors for main effect of drug and interactions with drug < 3 for error). For participants in the task feedback conditions (black and maroon filled symbols in Figure 2), levodopa tended to reduce reaction times in the instruction blocks, particularly for the larger aiming angles (angle x drug interaction, BF_incl_ = 2.169).

#### 3.1.2. Training improved performance across bins

In the subsequent training blocks, landmarks were removed, and the task-feedback manipulations were applied. Here, training sped-up reaction times across bins (see Figure 3) as shown by evidence for the main effect of training bin for reaction time (BF_incl_ = 1479.072). The effect of training bins on error and hit rates were subtler or absent, as shown by moderate evidence for a main effect of bin on error (error (BF_incl_ = 5.351) and moderate to strong evidence for excluding a main effect of bin for hit rate (BF_excl_ = 5039) and all interactions with bin (BF_excl_ > 7).

**Figure 3.**
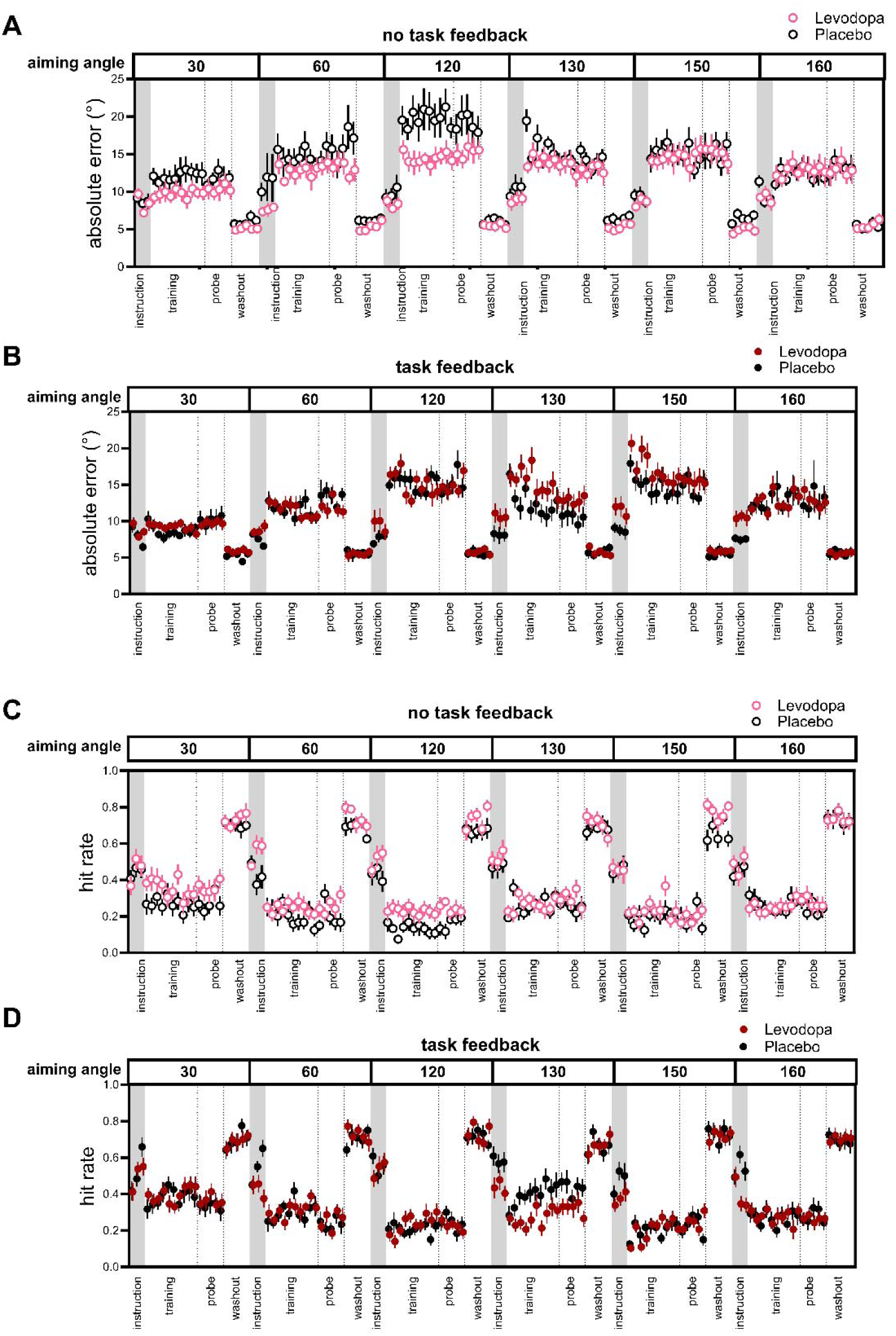
Bin-by-bin absolute errors for each of the aiming angles, for participants who completed training without task feedback (top panel) and with task feedback (bottom panels). Errors tended to be smaller and hit rate tended to be larger in the instruction blocks (gray regions in Figure 3), where participants aimed using landmarks, than in the subsequent training and probe blocks without aiming landmarks. At washout, participants rapidly reverted to aiming straight to the target, showing lower errors and higher hit rates than the preceding probe blocks.

To further examine effects of training bin, we analysed the influence of bin on reaction time and accuracy separately for each aiming angle. Reaction time decreased across bins for most angles, evidenced by strong main effects of bin (30°: BF_incl_ = 111.344; 60°: BF_incl_ = 89.126, 120°: BF_incl_ = 25.000; 160°: BF_incl_ = 82.324) or a strong bin × feedback interaction (150°: BF_incl_ = 33.693). The sole exception was the first angle set (130°), which showed evidence against a bin effect (BF_excl_ = 7.373). In contrast, accuracy exhibited strong evidence against bin effects at most angles (30°: BF_excl_ = 1240.124; 60°: BF_excl_ = 195.968; 120°: BF_excl_ = 310.699; 160°: BF_excl_ = 12.530), with exception of the initial angle set (130°: BF_incl_ = 534.048), which showed a reduction in errors across bins, concurrently with a maintenance of reaction times. Notably, even at angles showing robust changes in reaction time across bins (e.g., 30°, 60°, 120°, 160°), accuracy remained stable. We also did not find clear evidence that bin-by-bin training effects were modulated by drug condition or the combination of drug and feedback interactions: all bin × drug or bin x drug x feedback interactions showed evidence for exclusion across all angles for both reaction time and accuracy (all BF_incl_ < 3). These findings demonstrate that training-related performance changes across bins manifested primarily as reaction time improvements rather than accuracy changes, indicating that participants optimized response speed through practice without sacrificing accuracy.

#### 3.1.3. Non-linear effects of aiming angle on reaction times and errors

The effect of the aiming angles on reaction times and errors were non-linear for angles above 120° (see Figure 4). For example, with task feedback, errors were mostly similar for the 130°, 150° and 160° training blocks (as shown by weak evidence for difference between these angles, BF_10, U_ <2.5), Without task feedback, the 130° aiming direction resulted in overall lower error (BF_10, U_ =1.176e+10) and shorter reaction times than the 120° condition. We are unsure what drove this effect. One possibility is order effects. We used a fixed angle order across all conditions to prevent interactions between the drug condition and angle order. Across all participants, 130° was the first aiming angle, near the start of the session, whilst 120° was the last aiming angle, close to the end of a 2 hour task session. Aiming performance might be better early in the test session than late in the test session due to fatigue. Another explanation might be non-linear effects of aiming angle in visuomotor mental rotation tasks (Neely & Heath, 2010), for example, previous work showed faster reaction times when aiming at 210° than at 120° and 150° (Neely & Heath, 2010).

**Figure 4.**
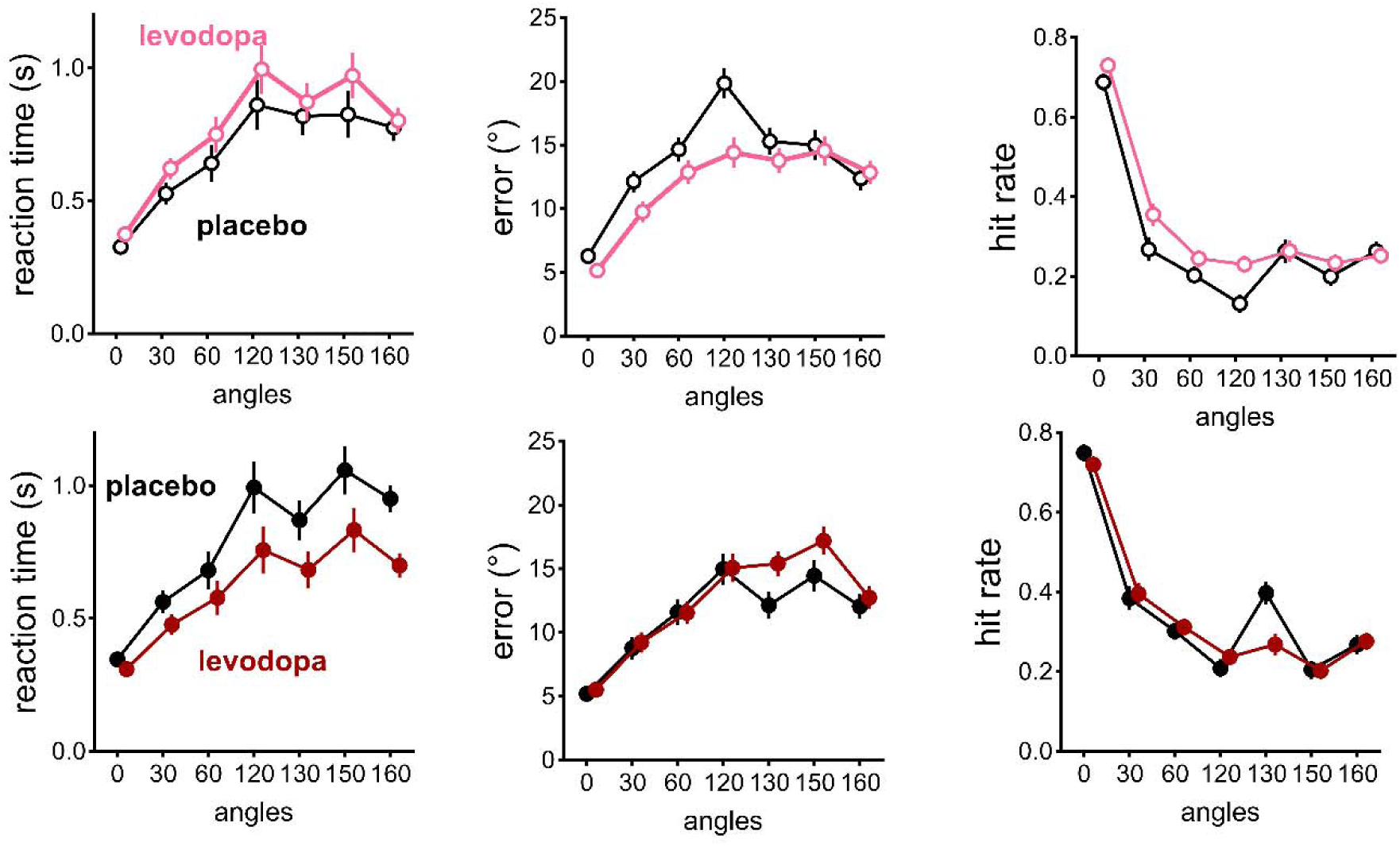
Without task feedback (top panels), levodopa increased the tendency to trade speed for accuracy, lengthening reaction times whilst reducing errors and increasing hit rates. Task feedback seemed to reduce this tendency to trade speed for accuracy (bottom panels): with task feedback, levodopa sped-up reaction times at a small cost to accuracy for larger angles, whilst mostly maintaining hit rates. This pattern held except for the 130° angle (which was at the start of the session), where the placebo group showed smaller errors and higher hit rates.

### 3.2. Dopamine altered performance both with and without task feedback

Dopamine differentially altered performance depending on the presence of task feedback (see Figure 4), although the effects of dopamine depended on feedback condition and aiming angle, as shown by Angle x Drug x Feedback interactions for error (BF_incl_ = 94.411) and reaction time (BF_incl_ = 6.71e+9); and Angle x Drug interactions for hit rate (BF_incl_ = 2.221 e+7).

In the absence of task feedback, levodopa (relative to placebo) augmented the tendency to trade speed for accuracy, resulting in overall longer reaction times (BF_10_, _U_ = 392303), smaller errors (BF_10_, _U_ = 10549), and higher hit rates (BF_10, U_ = 50741). Indeed, levodopa tended to reduce errors even when aiming straight at the target (main effect of drug, BF_incl_ = 2.105, BF_10, U_ = 14159). This effect of levodopa in trading-off speed for accuracy persisted into the post-training probe block, with moderate to strong evidence that dopamine improved accuracy (BF_10_, _U_ = 5.148) and hit rates (BF_10_, _U_ = 399.473) and lengthened reaction times (BF_10, U_ = 27.301).

In the presence of task feedback, dopamine sped up reaction times whilst maintaining achievement of the task goal (hitting the target), at a small cost of reduced aiming accuracy. Specifically, while levodopa (relative to placebo) had no effect on hit rates (moderate evidence for the null hypothesis for the main effect of Drug BF_01, U_ = 7.451), levodopa shortened reaction times and increased errors, particularly for the larger aiming angles, as shown by an Angle x Drug interaction for reaction times (BF_incl_ = 6.53e+16) and error (BF_incl_ = 29691). For the largest aiming angles (130°, 150°, 160°), these faster reaction times under levodopa conditions were accompanied by a small overall mean increase in errors of 2.584° (see Figure 4 bottom panel). Follow-up Angle x Bin ANOVAs tests for these three large angles yielded strong evidence for the effect of drug, particularly for the [Angle x Drug x Feedback interaction, BF_10, U_ = 2413], [BF_incl_ = 0.597]. Effects persisted into the post-training probe block where the task feedback was removed, as dopamine sped up reaction times (BF_10, U_ = 3.834e+17) whilst maintaining accuracy (strong evidence for the null hypothesis, BF_01, U_ = 13.469) and hit rates (moderate evidence for the null hypothesis, BF_01, U_ = 7.792).

### 3.2. Persistent effects of dopamine at washout

After the probe blocks, participants completed washout blocks when participants were instructed to stop re-aiming, and aim straight to the target. Figure 3 and 4 show that reaction times and errors dropped markedly, as re-aiming requires longer time for movement planning and is associated with more errors. Effects of levodopa in these washout blocks depended on the feedback condition and the aiming angle experienced in the previous training block, (Drug x Feedback interaction for reaction time, BF_incl_ = 4.625, Angle x Drug x Feedback interaction for error, BF_incl_ = 14.615). In participants who trained with task feedback, Levodopa did not clearly alter washout block reaction times or errors (inclusion bayes factors for the main effect of drug < 3 for both reaction time and error). In participants who trained without task feedback, whilst we did not find a clear effect of Levodopa on reaction times at washout (main effect of Drug, BF_incl_ = 1.105), Levodopa tended to reduce errors at washout, see Figure 3A (Angle x Drug interaction, BF_incl_ = 33.377, post-hoc tests BF_10,U_ = 20596).

### 3.3. Levodopa renders performance resilient to detrimental effects of feedback absence

The Drug x Feedback x Angle interactions for the reaction time and error data from the training blocks reported in the earlier paragraphs might also have been driven by differential sensitivity to the feedback manipulations in the levodopa or the placebo groups. We thus compared how the feedback manipulation altered the levodopa and placebo groups (see Figure 5), and also ran Feedback x Angle ANOVAs, separately for the placebo and levodopa groups.

**Figure 5.**
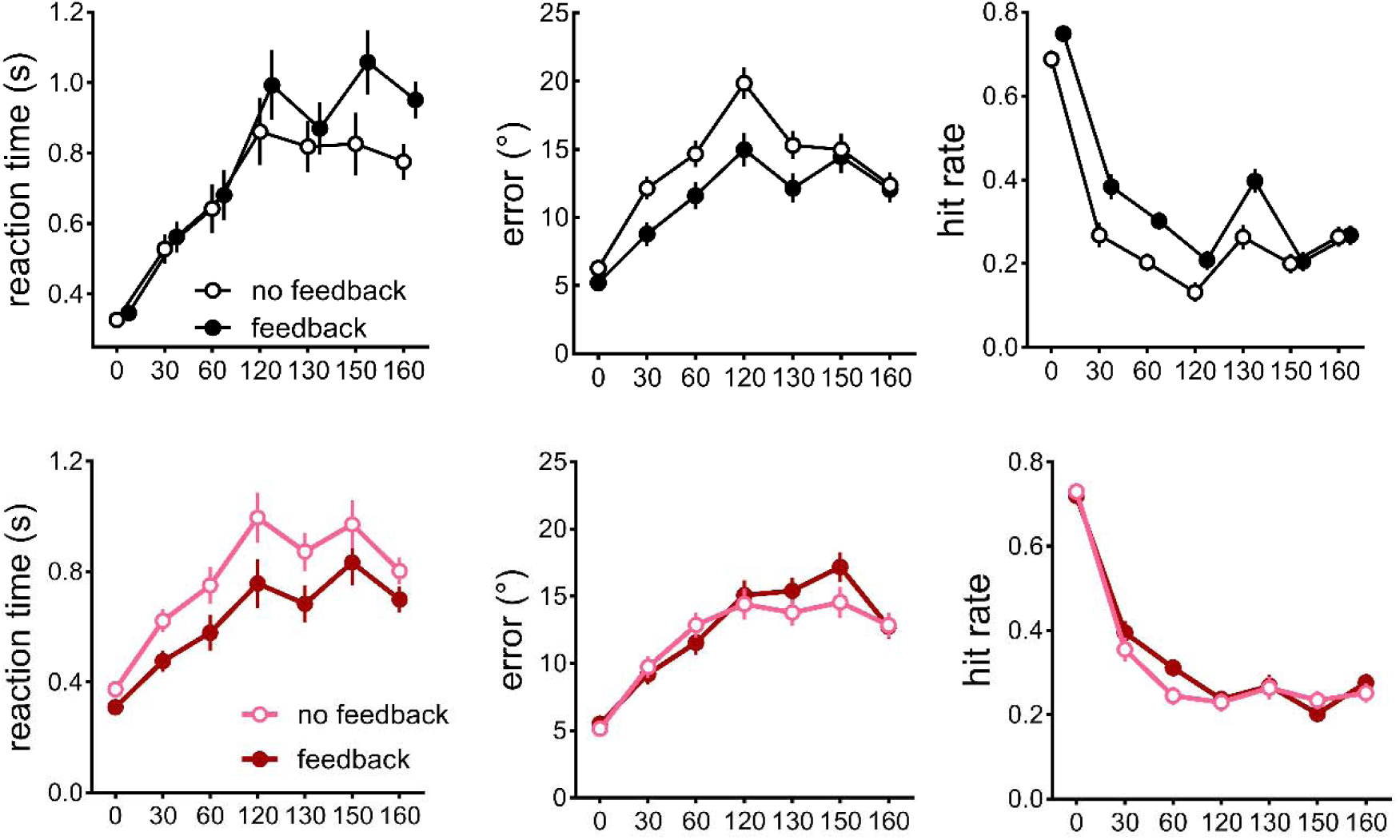
For the placebo groups (top panel), removing task feedback reduced hit rates, increased errors, and lengthened reaction times for the larger aiming angles (right panel). For the levodopa groups (bottom panel) hit rates and accuracy where comparable with/without task feedback (left/middle panels) in contrast to the placebo group (Figure 2). Also, in contrast to the placebo group, the levodopa group showed longer reaction times in the absence of feedback (right panel).

In the placebo groups, task feedback increased accuracy at the cost of slower reaction times for larger, more difficult aiming angles, as the task-feedback group showed higher hit rates (BF_10, U_ = 2.065e+13) and reduced errors (BF_10, U_ = 1.698e+8), (see Figure 4 top panels), and lengthier reaction times for larger aiming angles of 120° and above (BF_10, U_ = 4.491e+13).

In the levodopa groups), task success was not reduced by removing task feedback (see Figure 5 bottom panels), moderate evidence for null effect of task feedback for hit rate (BF_incl_ = 5.376), inconclusive effect of task feedback for error (BF_incl_ = 1.136). This maintenance of accuracy despite an absence of task feedback co-occurred with lengthened reaction times (angle x feedback interaction, BF_incl_ = 24.036, effect of feedback: BF_10, U_ = 4.219e+21), suggesting that the Levodopa groups traded off speed in movement planning for accuracy in the absence of task feedback (Figure 5 bottom right panel).

### 3.4. Feedback manipulations at training resulted in persistent post-training effects

After the training block, all participants were asked to re-aim in post-training probe blocks without any feedback manipulations. We assessed how previous training with/without feedback altered subsequent aiming performance in the probe block. Probe block aiming performance in the placebo groups depended on feedback presence at training: placebo groups who trained without feedback showed fewer hits (BF_10, U_ = 768705) and larger errors (BF_10, U_ = 80865484477), and shorter reaction times (BF_10, U_ = 2.81e+19) than their counterparts who trained with feedback. Thus, for the placebo groups, post-training performance was impaired by the absence of feedback at training. In contrast, the levodopa groups appeared to maintain the accuracy of their aiming regardless of whether they trained with or without feedback. Levodopa groups who trained without feedback showed similar hits and accuracy (moderate-strong evidence for the null hypothesis, hits: BF_01, U_ = 5.263; errors: BF_01, U_ = 11.848) but lengthier reaction times (BF_01, U_= 9.37) at post-training probe trials than their counterparts who trained with feedback. Thus, the Levodopa group participants maintained accuracy despite prior training without feedback, via a strategy of trading speed for accuracy.

### 3.5. Control measurements

Pharmacological manipulations of dopamine, particularly at larger doses than used here, can elicit undesired side effects such as nausea (for example, see Thirugnanasambandam *et al*., 2011; Chen *et al*., 2020). However, none of our participants reported this. Exogenous dopamine can also change mood state (Vo *et al*., 2018), however, here, we did not find evidence that levodopa altered our participants mood state as indicated by inconclusive evidence for a change in mood state on the Bond Larder Scale (BF_excl_ = 1.288). Blood pressure and heart rate was also unaltered, as Time (before drug administration, 30 minutes after drug administration, 2 hours after drug administration) x Drug ANOVAs yielded moderate evidence for the null hypothesis for Time x Drug interactions for blood pressure (diastole: BF_excl_ = 4.149, systole: BF_excl_ = 8.547) and mood (BF_excl_ = 3.681), and heart rate (BF_excl_ = 3.731). In addition, participants were not above chance at accurately guessing the drug condition, as shown by Bayesian binomial tests (BF0+ = 4.685). Neither experimenters nor participants were above chance at accurately guessing the drug condition, as shown by moderate evidence for the null hypothesis from Bayesian binomial tests (participant: BF0+ =6.181, experimenter: BF0+ = 4.603).

### 3.6. Exploratory analyses

Given the hypothesized role of dopamine in vigour, it is plausible that dopamine might also alter indices movement vigour, such as peak velocity. Thus, exploratory analyses on effects of levodopa and feedback manipulations were run. When landmarks were provided for aiming in the instruction block, Levodopa did not alter peak velocity (main effect of Drug, BF_incl_ = 0.834, Angle x Drug interaction, BF_excl_ = 6.623). However, Levodopa reduced peak velocity when landmarks were removed in both the training and probe blocks, particularly for the larger aiming angles, as shown by Angle x Drug interactions during training (BF_incl_ = 644.636) and probe (BF_incl_ = 21230). Once the requirement for aiming was removed in the washout blocks, Levodopa ceased to alter peak velocities (evidence for no main effect of drug, BF_excl_ = 2.736, BF_excl_ = angle x drug interaction= 68.824). Thus, levodopa primarily altered peak velocity when tasked with aiming away from the target by specified angles. There was no general effect of levodopa on peak velocity, consistent with (Palidis *et al*., 2021).

Task feedback was associated with lower peak velocity when aiming, as shown by a main effect of feedback for at landmark (BF_incl_ = 2.834) training (BF_incl_ = 3.834), and probe block (BF_incl_ = 7.033), as well as after participants were instructed to stop re-aiming in the washout block (BF_incl_ = 6.636). The presence of the effect of task feedback in blocks without the task feedback manipulation (the landmark, probe, and washout blocks) suggests that recent experience of task feedback in the preceding or subsequent training blocks tends to slow peak velocities. This slowing of peak velocities with task feedback is similar the effect of task feedback in slowing reaction times described above.

## Discussion

Here, in a preregistered, double-blind, placebo-controlled study, we examined how task feedback altered learning and performance in a task that where participants learn to aim away by various angles relative to the presented target, which is a key explicit strategy used in the classic visuomotor rotation task. Furthermore, we also tested whether the effect of task feedback on this task depended on dopamine.

Participants were either provided with salient information on their task success/failure, or deprived of task success information about their performance. In a task that prioritised accuracy, not speed, exogenous dopamine appeared to help participants optimise task success. When deprived of task success information: levodopa increased the number of target-hits and increased accuracy: and this was achieved by participants varying the speed accuracy trade-off, as shown by an increase in reaction times. When feedback about task success was provided, levodopa sped-up reaction times whilst maintaining overall task success (target-hits). Our results provide direct evidence for a role of dopamine in distinct processes that shape performance in explicit sensorimotor adaptation: one driven by internally generated estimations of performance, and another driven by external information about performance.

To the best of our knowledge, our study is the first to provide causal evidence for a role of dopamine in motor performance when motor learning was driven by internally generated estimations of performance in neurotypical humans. We built upon our previous study (Leow *et al*., 2024), which provided extrinsic task feedback information throughout the task. In that work, we showed that increasing dopamine availability improved task performance by increasing a propensity to better adhere to task goals, suggesting engagement of instrumental motivation to optimize adherence to task goals (Leow *et al*., 2024). However, because task feedback was always present, it was unclear whether the effects of dopamine in that study resulted from effects of dopamine on intrinsic motivational processes that operates in the absence of task feedback, and/or extrinsic motivational processes that rely on task feedback. Here, by explicitly manipulating the presence and absence of task feedback, we provide direct evidence for a role of dopamine in process(es) driven by internally generated estimations of performance. Whilst neuroimaging evidence indirectly implicates the striatal mesolimbic areas when learning in the absence of extrinsic feedback (Daniel & Pollmann, 2012; Guggenmos *et al*., 2016; Ripolles *et al*., 2016), and pharmacological manipulations of dopamine show a role of dopamine in modulating motor or cognitive task performance in exchange for rewards (De Vries *et al*., 2010; Chong *et al*., 2015; McGuigan *et al*., 2019; Michely *et al*., 2020), there is a surprising dearth of direct evidence for a role of dopamine in motor performance in the *absence* of extrinsic task or reward feedback. Whilst one previous study showed that modulating dopamine availability via levodopa or a dopamine antagonist altered language learning in the absence of external task feedback (Ripollés *et al*., 2018), that study lacked a placebo control, and thus it is unclear if effects were driven by levodopa improving learning or by the dopamine antagonist impairing learning. Here, in our placebo-controlled design, we show that exogenous dopamine improved motor performance, both in the absence of task feedback, and in the presence of task feedback. By directly contrasting effects of dopamine on a design utilizing only minimal differences between conditions with and without feedback about task success, we demonstrated separable effects of levodopa on performance with and without task feedback.

Exactly how did levodopa alter performance with and without task feedback in our study? Compared to the placebo group, participants in the levodopa group seemed better able to leverage task feedback to optimise performance, speeding-up reaction times without incurring a cost to accuracy for the smaller aiming angles, although there was a small drop in accuracy for the larger aiming angles that mostly maintained hit rates. The finding that exogenous dopamine improves feedback-based performance is consistent with previous findings of improved motor and cognitive task performance with levodopa in the presence of feedback (De Vries *et al*., 2010; Chowdhury *et al*., 2013; Shellshear *et al*., 2015). This effect seems most prominent with larger, more difficult aiming angles, which may partly explain why previous work using subtler manipulations do not show effects of dopamine manipulations (Quattrocchi *et al*., 2018; Palidis *et al*., 2021). The picture differed in the absence of task feedback. Whilst accuracy suffered in the absence of feedback about task success in the placebo group (Schaefer *et al*., 2012; Kim *et al*., 2018; Leow *et al*., 2018; 2020; Oza *et al*., 2020; Tsay *et al*., 2022), exogenous dopamine here appeared to help maintain accurate performance, such that accuracy was similar regardless of whether feedback about task success was present. This improved capacity to maintain task performance seems consistent with an increasing body of work showing that exogenous dopamine tends to blunt the effect of manipulations of task-related information on performance, such as the valence of potential task outcomes (e.g.,Guitart-Masip *et al*., 2014; Vellani *et al*., 2020). In summary, our findings suggest that dopamine influences motivation, both to improve internally determined task performance, and to improve performance shaped by extrinsic task feedback.

How might levodopa have changed motor performance in the absence of extrinsic information about task success during this task? Levodopa is a precursor to dopamine, which increases brain dopamine availability. Animal studies show that levodopa increases phasic firing of striatal dopamine cells (Willuhn *et al*., 2014). We speculate that levodopa increased the amplitude of dopamine signalling in response to internal evaluations of task performance. This interpretation is in line with recent findings in animals which demonstrate a key role of dopamine in internal evaluations of performance (Guggenmos *et al*., 2016; Duffy *et al*., 2022). For example, spontaneous dopamine activity in songbirds correlate with self-evaluation of performance quality in the absence of external rewards, cues, or perturbation (Duffy *et al*., 2022). Similarly, Markowitz *et al*. (2023) showed that in rats acting in the absence of explicit reward (i.e., in darkness), dopamine release was followed by an increase in spontaneous actions, and that increased dopamine release after such actions was associated with an increased probability of repeating such actions. In a similar vein, one fMRI study in humans has suggested that activation of the striatum in response to reward, traditionally interpreted as a neural representation of the value of that reward, might instead be better interpreted as a representation of how well those rewards align with one’s internal representation of a task goal (Frömer *et al*., 2019). In a task where participants were asked to choose the best option or the worse option, neural activity in value-related regions were better accounted for by how well the reward aligned with the task goal, rather than how rewarding each option was (Frömer *et al*., 2019). These results, as well as ours, are broadly supportive of the view that dopamine in the striatum is instrumental to internal estimations of task performance relative to some goal (Ptasczynski *et al*., 2022), consistent with the idea that dopamine has a generalised role in prediction (Gardner *et al*., 2018).

## Summary

The role of intrinsic motivation has long been recognised within the context of motor learning (Wulf & Lewthwaite, 2016), and theories have broadly implicated brain dopamine in driving intrinsic motivation during skill learning (Lewthwaite & Wulf, 2017). However, to date, such theories have yet to be substantiated by evidence in healthy humans. Our findings provide this missing evidence. We show that exogenous dopamine alters performance during an explicit motor learning task, both in the absence and in the presence of extrinsic feedback, demonstrating a key role of dopamine in optimising motor performance during skill learning, even in the absence of external task feedback. Our study thus provides direct evidence for the role of dopamine in performance guided by internal estimations of performance in humans. This work not only has implications for implementation of motor learning in rehabilitation contexts in populations with changes in brain dopamine function (e.g., healthy ageing; (Bäckman *et al*., 2006), but will also improve future implementation of dopamine pharmacotherapy as an adjunct strategy to optimise motor rehabilitation (Scheidtmann et al., 2001; Rosser et al., 2008).

## Acknowledgements

We would like to acknowledge funding support from the Australian Research Council (DP230102179) and the Brain Foundation.

